# Low-cost arabinose induced genetic circuit for protein production: Modeling

**DOI:** 10.1101/2021.12.31.474657

**Authors:** Christian J. Flores Gómez, Jorge L. García Barrera, Edgar Valeria de la Cruz

## Abstract

A low-cost reagent-producing genetic circuit was designed during this work. Its functioning is based on a positive feedback loop induced by a small amount of arabinose, allowing users to obtain reactants in a safe, constant, and controlled manner. The “design only” approach to the project allows us to work in different kinds of computational models, thus, an ODE-based model was thoroughly developed and a cellular automata-based one was experimented with.

Working on the ODE model, equilibrium states and system stability were studied. Circuit properties were also focused on one of which was a high concentration of interest protein produced by low inductor inputs. As a result, a mathematical expression capable of describing the quantity of produced reagent was obtained. In addition, the cellular automata model offers a new perspective, given its differences from the ODE model e.g. this type of model is a stochastic analysis and describes each cell individually instead of describing the whole cellular population.

## I. Introduction

### A. Genetic circuits

Genetic circuits are electrical circuits analogs, used in Synthetic Biology. They are engineered from individual genes with various functions. This kind of circuit generally includes the following parts:

- A promoter, functioning as the On/Off control for the circuit
- A Ribosome Binding Site (RBS), where the ribosome attaches to initiate transcription of the protein
- Coding Sequence (CDS), containing the information about our target protein
- Terminator sequence, marking the end of the target protein and causing transcription to stop

### B. Mathematical models

Mathematical models are descriptions of reality with predictive capacity. Traditionally differential equations have served as means to explain an unlimited number of phenomena. However, some generate too complex equations without existing analytical solution. For these cases, mainly studied from nonlinear dynamics, techniques have been developed that allow them to be studied successfully. These techniques range from a reduction in the number of constants to their linearization for the study of their equilibrium and stability. Positive feedback genetic circuits have non-linear dynamics that will be studied in this work for our specific genetic circuit.

### C. Cellular Automata

The incredible complexity of biological systems usually leads to mathematical models composed of non-linear differential equations (NDE’s). However, an NDE-based model aiming to describe in a realistic manner this type of system presents a few challenges. Initially, the structure of the system may be unknown or still be under research. Secondly, when trying to solve the NDE’s, computers with a high processing speed and copious amounts of memory are needed. Finally, a biological system is composed of a large number of intricate and interconnected variables affecting each other.

Cellular automates make ideal structures to design complex computational systems thanks to their capacity to receive properties describing the interactions between elements and the behaviors leading to them. The complexity of modeling bacteria as a system is linked to protein synthesis and genome reading. In synthetic biology, these phenomena become modifiable systems whose behavior can be altered in a specific way by reprogramming the genetic code. For that reason, analyzing the dynamics of interaction and protein production using cellular automata is very simple.

### D. Functioning

The circuit (see fig.1) is designed to be simple in use, reduce costs and provide control in protein production. The circuit is activated with a simple sugar, arabinose, this molecule was chosen mainly for its relatively low cost compared with other commonly used inductor molecules (e.g., IPTG). The low-cost arabinose induced genetic circuit is a positive feedback loop activable with a 5-carbon sugar commonly used as a non-gratuitous inductor, meaning this sugar can not be metabolized by the microorganism in which is introduced (*Escherichia coli*. in this case). Once the circuit is activated once it’ll keep turning itself “ON”, producing proteins until the resources available deplete. The *in silico* demonstration of the low arabinose use in this circuit is one of the goals the work here. Positive feedback loop circuits have been proven as the foundation for biestabilty[7]; the practical applications for biestability are countless, in this case, the presence of biestability means that, even if there’s no sugar present in the media and *E. coli* has degraded all the arabinose in it, the circuit can maintain the production of protein. This is not possible in conventional circuits.

**Fig. 1.**
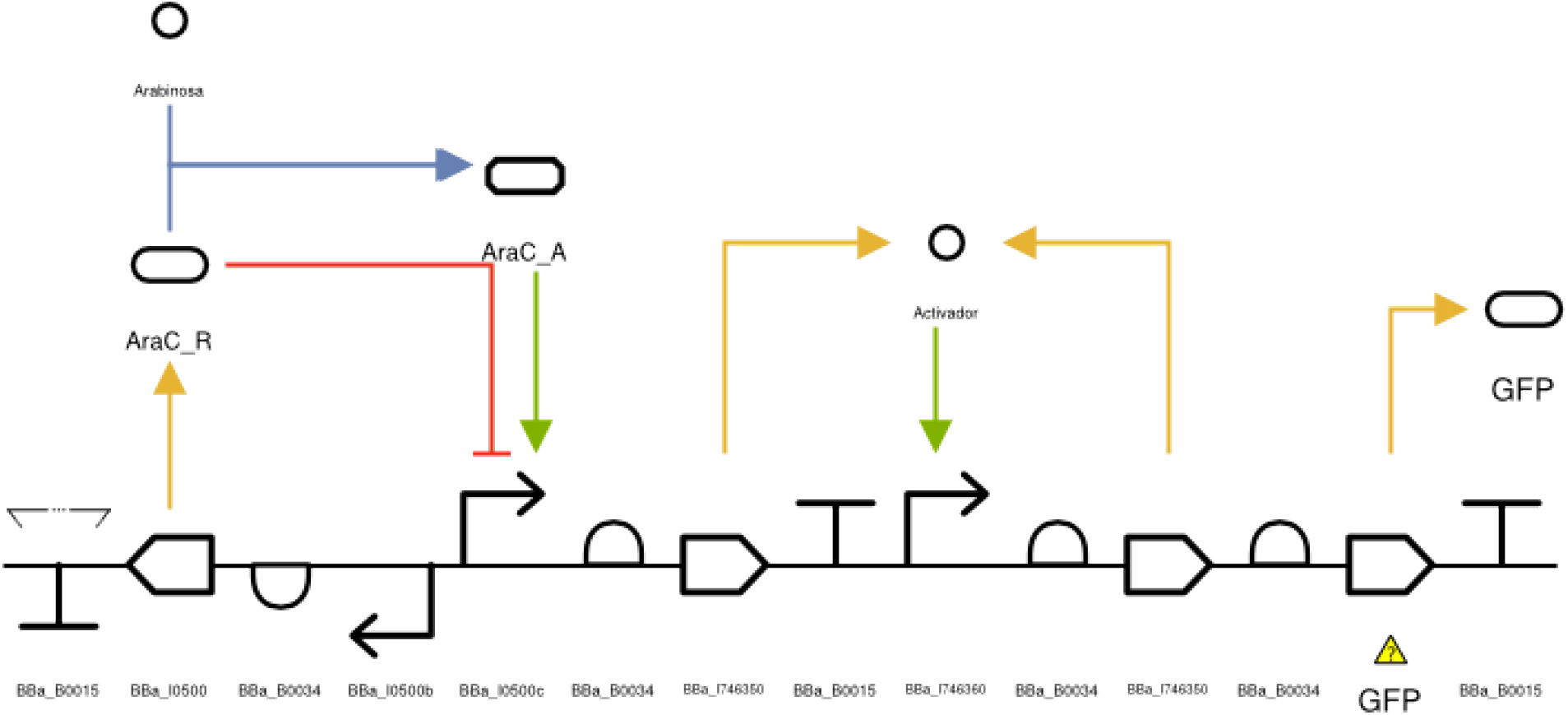
Graphic representation of the genetic circuit.

## II. Mathematical model

### A. Chemical equations

With the simplification of the differential equations that will describe the system dynamics as a goal, some hypothesis will be assumed. The approach here used to propose the differential and chemical equations is explained in [8]. Here, the promotor is assumed as a representation of the whole transcriptional unit. Transcription and translation are treated as a single step, this implies that mRNA and ribosomes are not present in this model. Given the RNA polymerase concentration is usually a lot higher than that of the plasmid containing the circuit, it is assumed its concentration does not change based on the interactions with the circuit. Thus, RNA polymerase does not appear in this model. Varitations in concentration by cell division are also not accounted for.

Let be the circuit shown in figure 1, assuming enough time has passed (time *τ*_1_), AraC protein is producing constitutively and has reached an equilibrium concentration and thus it is constant. It is also assumed enough time has passed (*τ*_2_) for araC to diffuse into the medium and joins to the P1 promoter acting as a repressor. Once time *τ*_1_ + *τ*_2_ has passed, it is assumed arabinose has been added. Knowing araC protein maintains itself attached to the promoter forming a complex in presence or not of arabinose [9], gives us reasons to consider the concentration of P1 promoter not attached to araC as negligible. According to [12] arabinose can associate to araC even if it is already forming a complex with the promoter, meaning, it is not necessary for araC to be free to associate with arabinose. These ideas are resumed in the following chemical equation:^1^

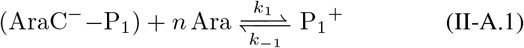

Where (AraC^−^−P_1_) is the promoter-repressor complex, Ara is arabinose and P_1_^+^ is the promoter-activator complex (an activated promoter). This promoter can attract the RNA polymerase, initiating transcription and then translation of activator A. This promoter is also an inexhaustible reagent:

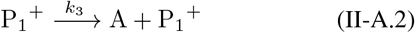

The activator protein A can attach itself to promoter P_2_, activating it.

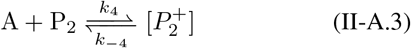

now, this activated promoter 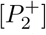 can attract RNA polymerase, start transcription and then translation of the activator A and the protein of interest, in this case GFP and as in the previous case, it is an inexhaustible reagent

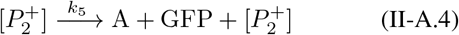

Basal expression is a process to keep in mind in this model and it is considered in the following equation.

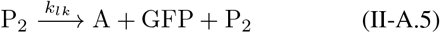

Lastly, activator A and target protein GFP degradation were also considered due to physical and biological causes.

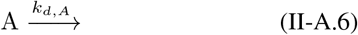

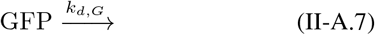

### B. Differential equations

Once the chemical equations have been presented a differential equation model based on law of mass action can be constructed. It can be observed that *k*_5_ *>> k*_*lk*_(see equations II-A.4 y II-A.5), meaning basal transcription rate is much lower than activator mediated transcription. This is valid for positively regulated promotors. Now, be [*P*_1,*T*_] the total concentration of promoter 1, and with the previous arguments:

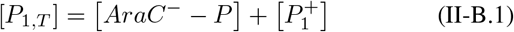

Now 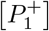 dynamics Is given by^2^

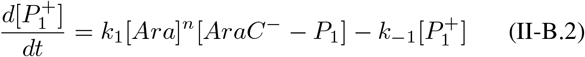

Some time after adding arabinose is reasonable to think that 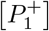 will arrive at a constant equilibrium concentration, i.e., 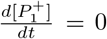.Using equation II-B.1 and rearranging it the following equality is obtained:

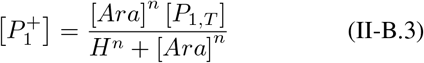

Where

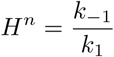

*H* being the Hill constant and *n* the Hill coeficent. Through simplifications and assumed hypothesis a Hill function is capable of modelling P_1_^+^ concentration based off Ara concentration was obtained. This relation is the key to the one between GFP concentration and Ara. In figure 2 a graphical representation can be seen, the parameters were chosen arbitrarily. Now the complete dynamics of the circuit can be modelled. The dynamic of the Promoter 2 attached to activator protein 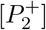, activator protein [*A*] and target protein [*GFP*] are of special interest

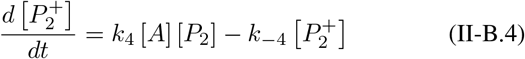

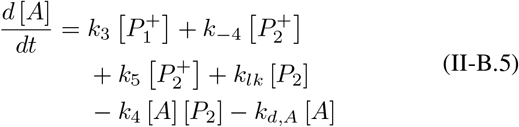

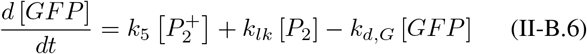

**Fig. 2.**
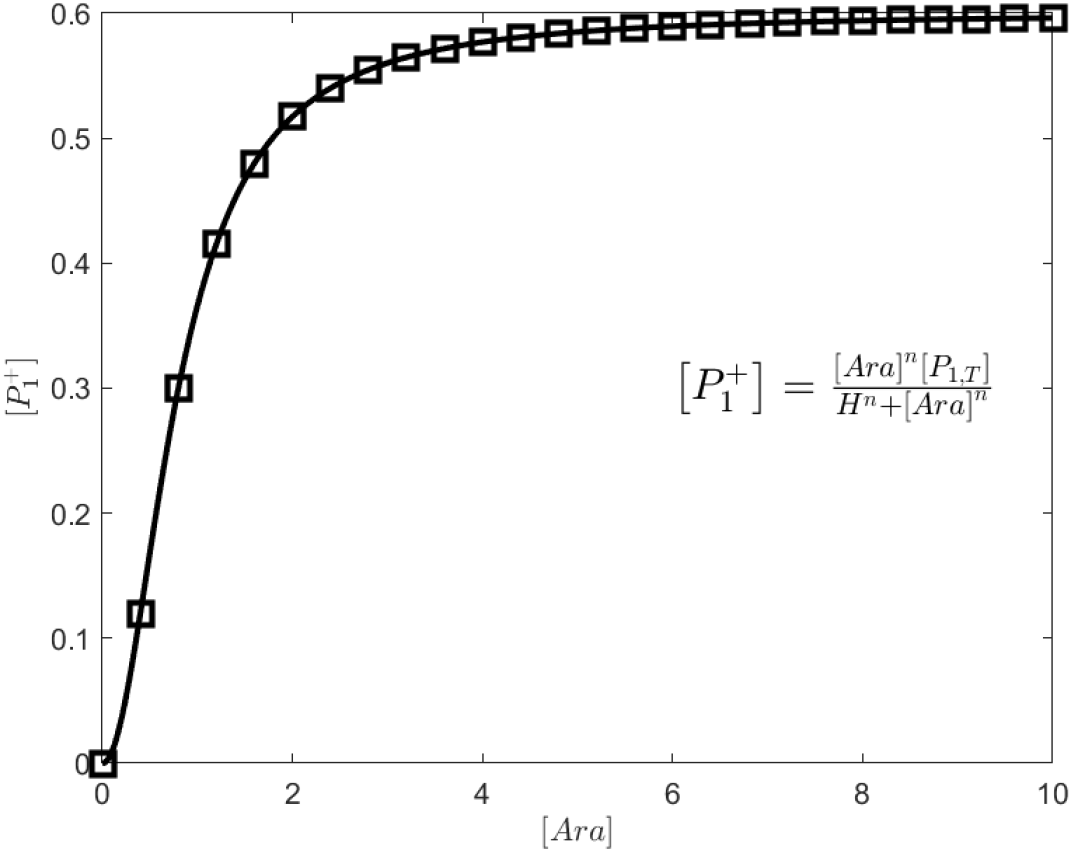
The 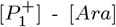 dependency is described via a Hill function. The following parameters were chosen 0 ≤ [*Ara*] ≤ 10; *n* = 2; [*P*_1,*T*_] = 0.6; *H*^*n*^ = 0.8.

Equations II-B.4 y II-B.5 are coupled and thus their analysis is fundamental to understand the circuit dynamics. The total concentration of promoter 2 [*P*_2,*T*_] is constant and is given by

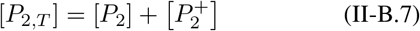

Substituting equations II-B.4 y II-B.5 and rearranging

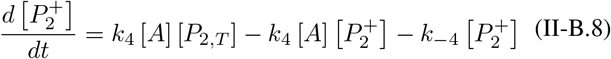

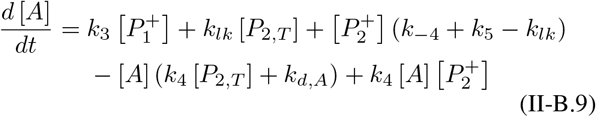

In dimensionless form^3^

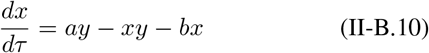

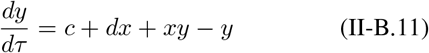

Where^4^

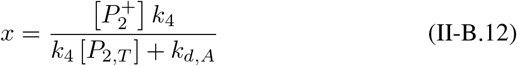

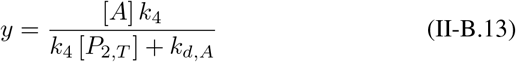

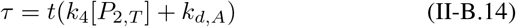

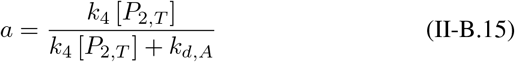

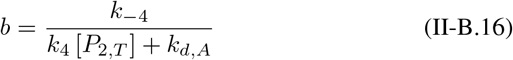

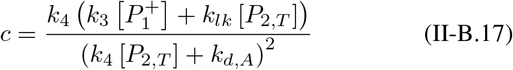

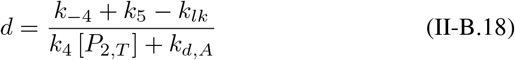

If *k*_1_, *k*_−1_, *k*_2_, …, *k*_*d,G*_, [*P*_2,*T*_] *>* 0, ergo, all the constants in the model are positive, the following properties are obtained

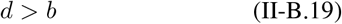

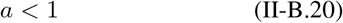

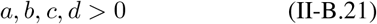

II-B.19 It’s obtained from *k*_5_ ≫ *k*_*lk*_ and II-B.20 from *k*_*d,A*_ *>* 0.

### C. Equilibrium and system stability

The system is in equilibrium if^5^

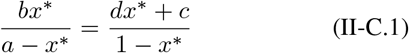

Reorganizing following solution for *x*^*^ is obtained

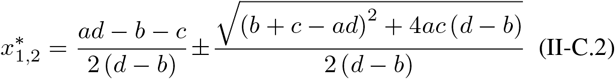

As *a, b, c, d* > 0 y *d* > *b*, then

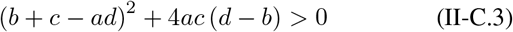

Therefore 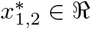 such that 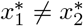.

The system will always have 2 real equilibrium states. There is no bifurcation..

As 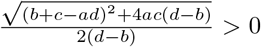, the sign that *x*^*^ takes depends on 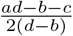.

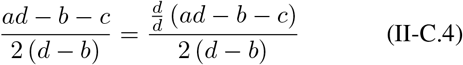

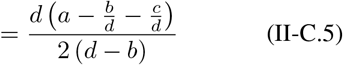

Being 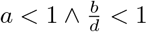 we can approximate

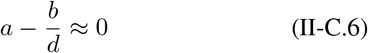

Thus

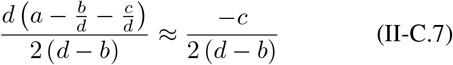

Using the same approximation, we get the following result.

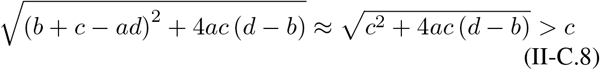

Therefore, if the approximation is valid, it can be assured there are always a positive and a negative state of equilibrium. Only one of this two has biological interpretation. Based on the next equality

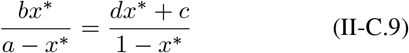

The positive equilibrium must satisfy *x*^*^ *< a <* 1.This will give a maximum value to which the system cannot arrive.

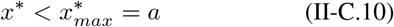

The system equilibrium stability is also studied. And thus, linearization of the differential equation system is necessary. This is done calculating the differential equation system’s Jacobian. The Jacobian of the system *J* is given by

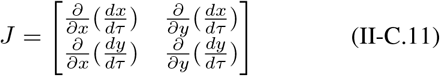

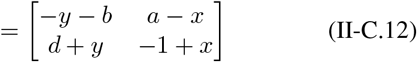

Be *T* the matrix trace and Δ its determinant

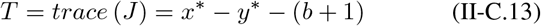

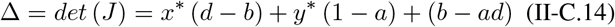

Using the same approximation 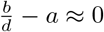, the following is obtained

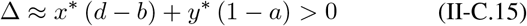

Now

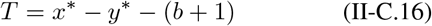

Substituting 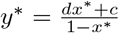

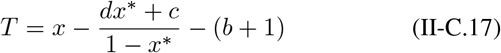

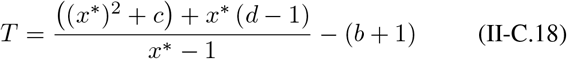

*T <* 0 can be proven for all *a, b, c, d, x*^*^ *>* 0 and *x*^*^ *<* 1. As long as the used approximation used is valid and under the proposed hypothesis, the system will reach a unique stable equilibrium. Figure 3 illustrates a phase plane of the system.

**Fig. 3.**
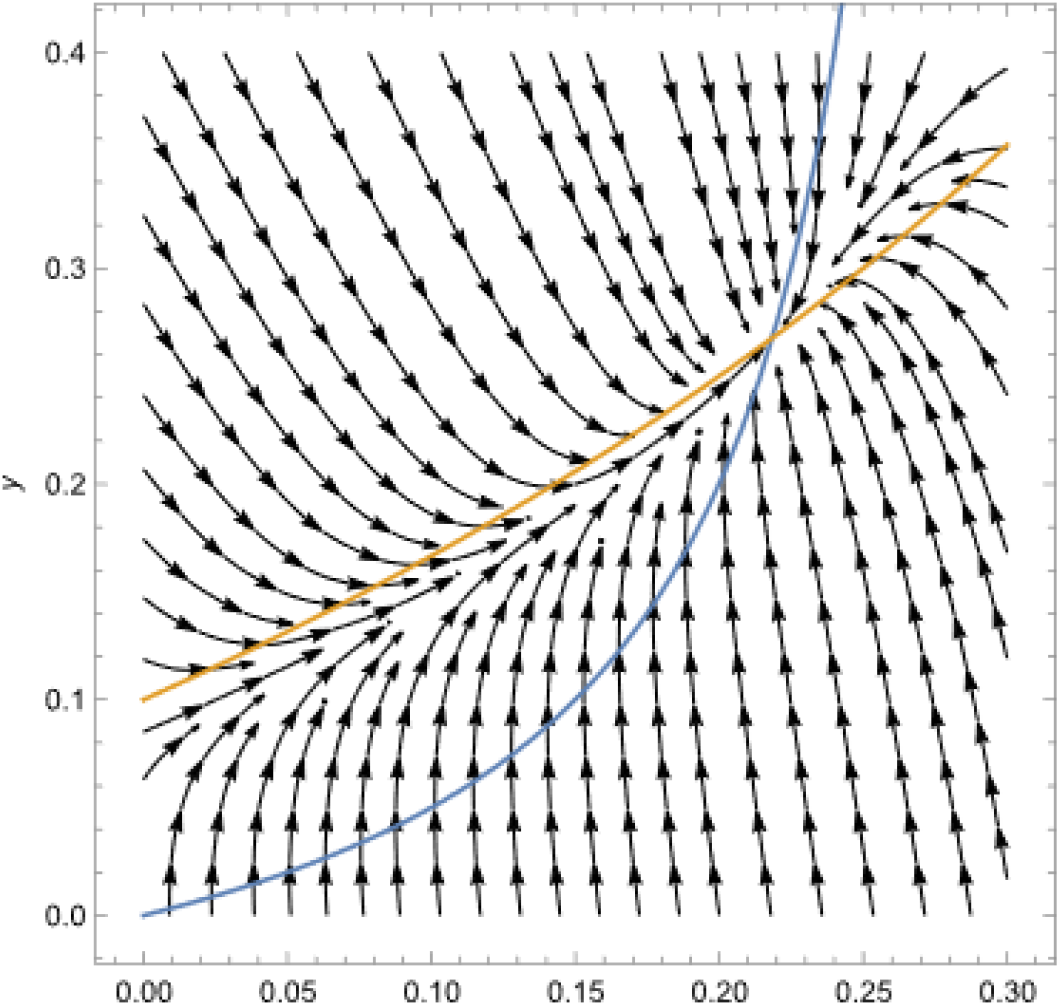
Phase plane of the system with *a* = 0.3, *b* = 0.1, *c* = 0.1, *d* = 0.5.

### D. Differential equation numeric solution

Next, numerical solutions for the differential equations system in various initial conditions are graphed(see fig. 4, 5 y 6). Ode45 solver from MATLAB®was used to solve them. Parameters and units were chosen arbitrarily:*k*_4_ = 0.8; *k*_−4_ = 0.4; *k*_3_ = 8; [*P*_1,*T*_] = 8; [*Ara*] = 1000; *H* = 0.2; *n* = 2; *k*_*lk*_ = 0; [*P*_2,*T*_] = 8; *k*_5_ = 3; *k*_*d,A*_ = 8; *k*_*d,G*_ = 0.4.

**Fig. 4.**
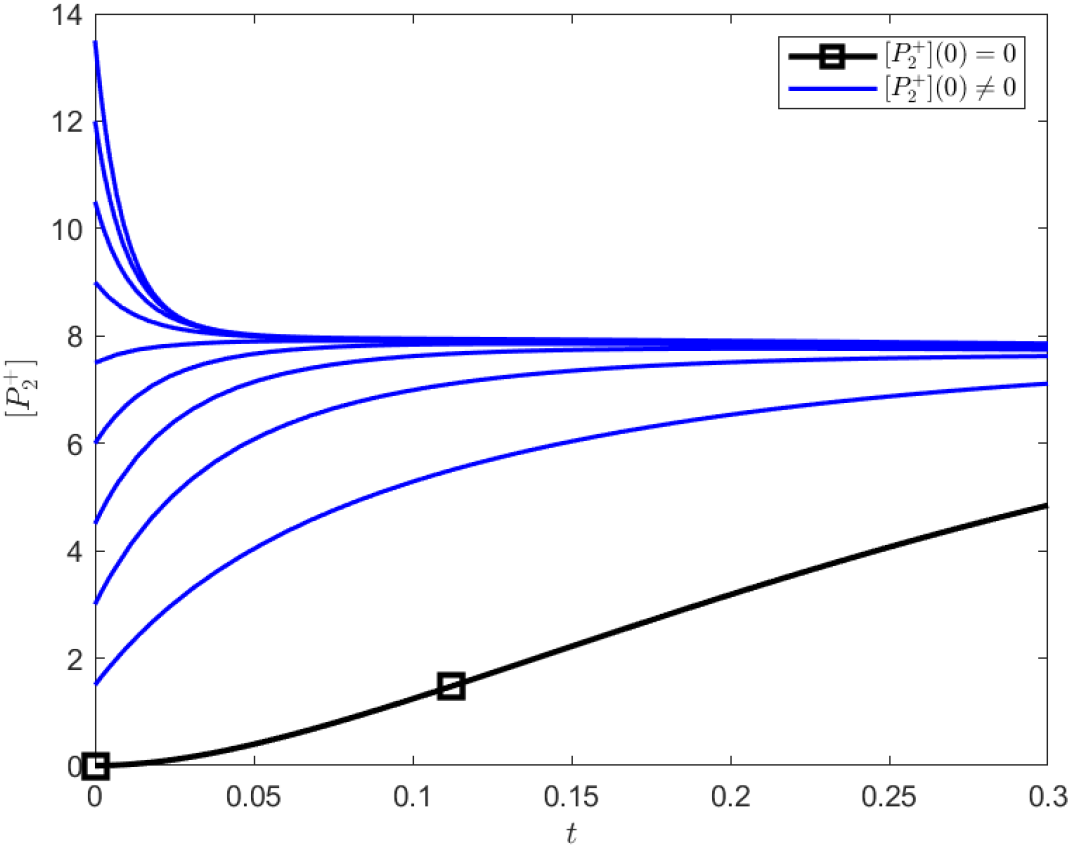
Activated promoter 2 concentration dependent of time with various initial conditions.

**Fig. 5.**
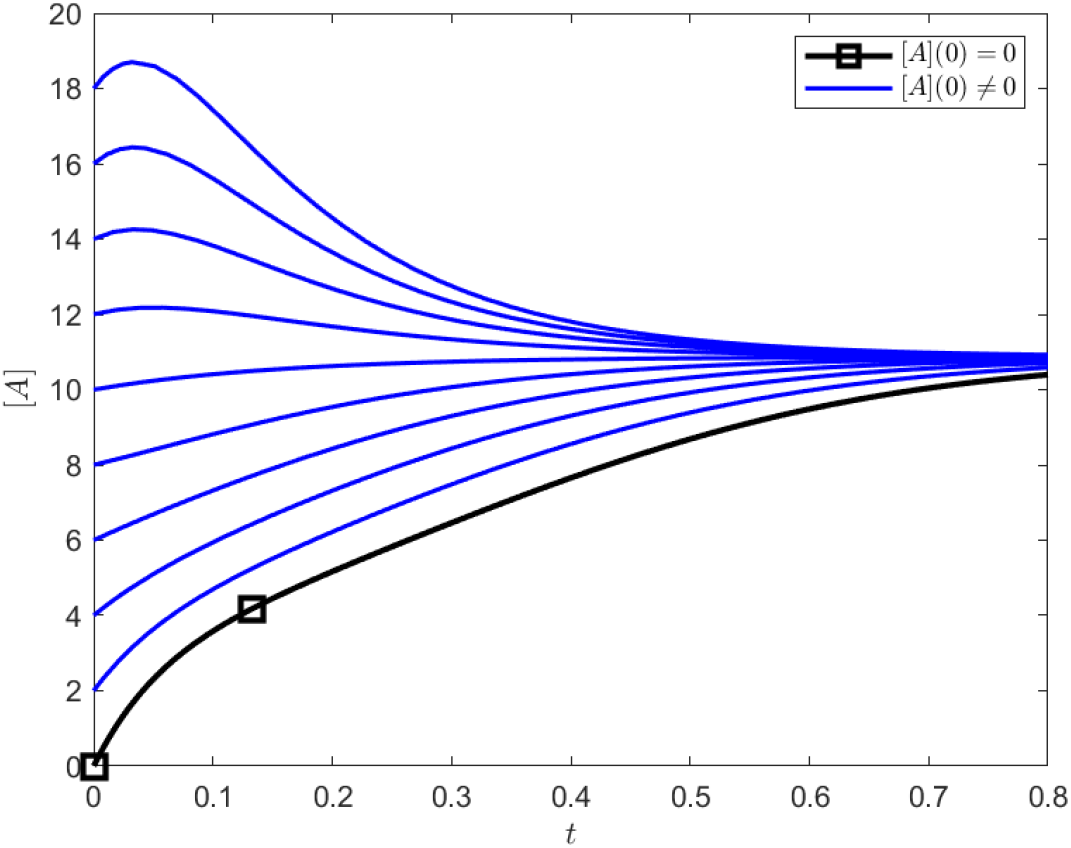
Activator protein concentration dependent of time with various initial conditions.

**Fig. 6.**
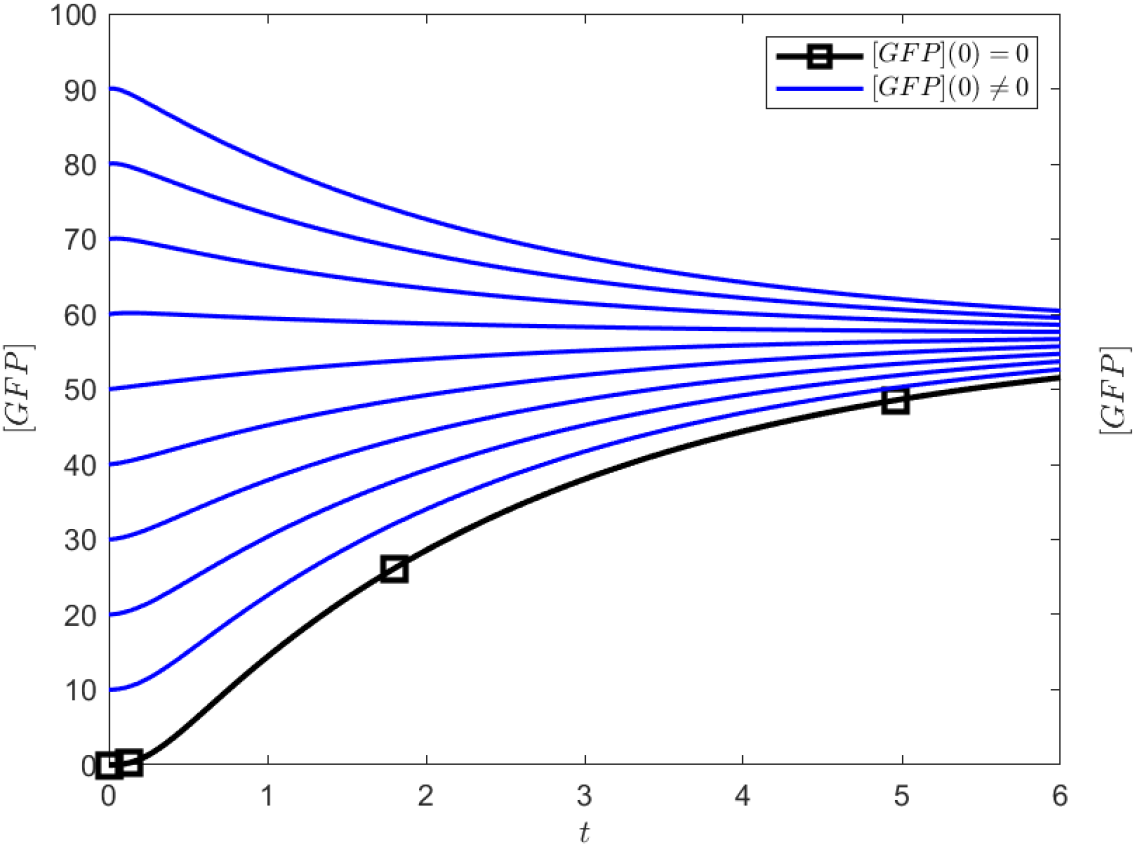
GFP protein concentration dependent of time with various initial conditions.

### E. Hysteresis

Memory or hysteresis is one of the main characteristics of this genetic circuit. Once the inductor is added, the circuit remembers its active state. The target protein will be produced even after degradation of the inductor sugar, keeping the circuit in active state. The differential equations system can model this behavior:

As 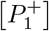 and *k*_*lk*_ equal to zero then *c* = 0. All the previous properties are still true: *a, b, d >* 0, *d > b, a <* 1. Substituting *c* = 0 on the *x*^*^ equilibrium solutions

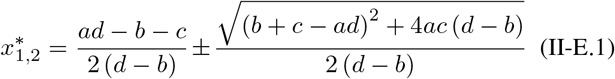

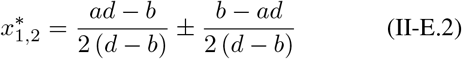

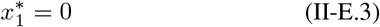

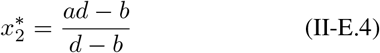

It is known that *d > b*, so for a big enough *a* we can assume *ad > b*. Then 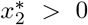. Two equilibriums with biological significance exist, a positive one and one equal to 0. Now the stability of these two equilibria is studied. When *x*^*^ = 0 and *y*^*^ = 0.

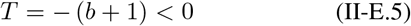

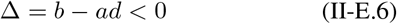

This means this equilibrium is a saddle point and thus unstable. When *x*^*^, *y*^*^ *>* 0

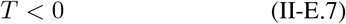

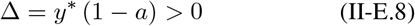

This implies equilibrium here is stable. The phase plane in 7 is an example. Two equilibriums can be observed (stable *x*^*^, *y*^*^ *>* 0 and unstable *x*^*^, *y*^*^ = 0).

While the basal production is negligible, and arabinose has no been added the system is in unstable equilibrium where the concentrations of 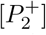, arabinose and GFP are 0. Nevertheless, a fluctuation in the system can take it to its stable equilibrium, where the concentrations are no longer zero. When the system is on a stable equilibrium state where the concentrations area greater than zero, if the arabinose degrades in its entirety the system will reach a new equilibrium state such that *x*^*^, *y*^*^ *>* 0. The system remembers its state, this phenomenon it’s called hysteresis. The hysteresis can be visualized with a (see fig. 8) obtained via numerical solutions and variation in the arabinose concentration.

**Fig. 7.**
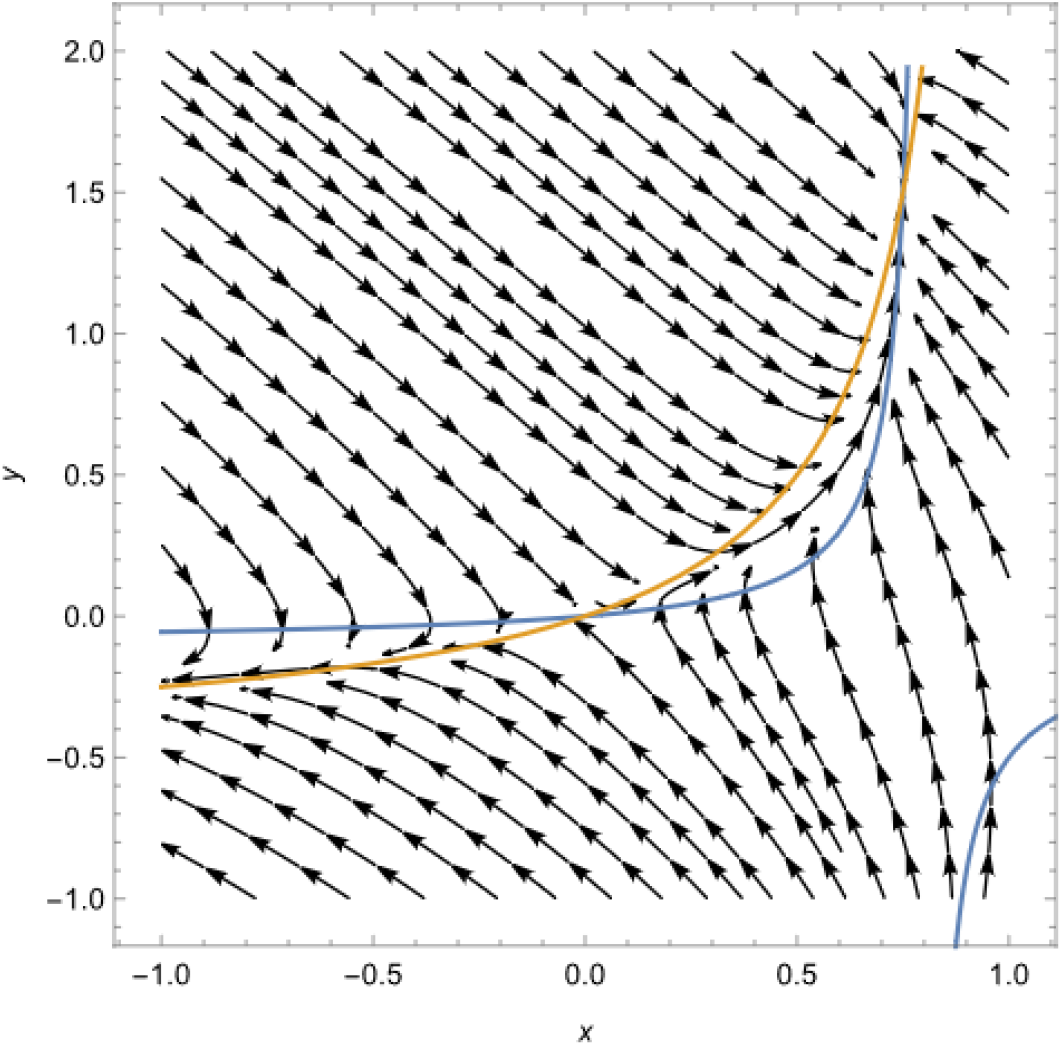
Plane with 2 equilibria. The values are *a* = 0.8, *b* = 0.1, *c* = 0, *d* = 0.5.

**Fig. 8.**
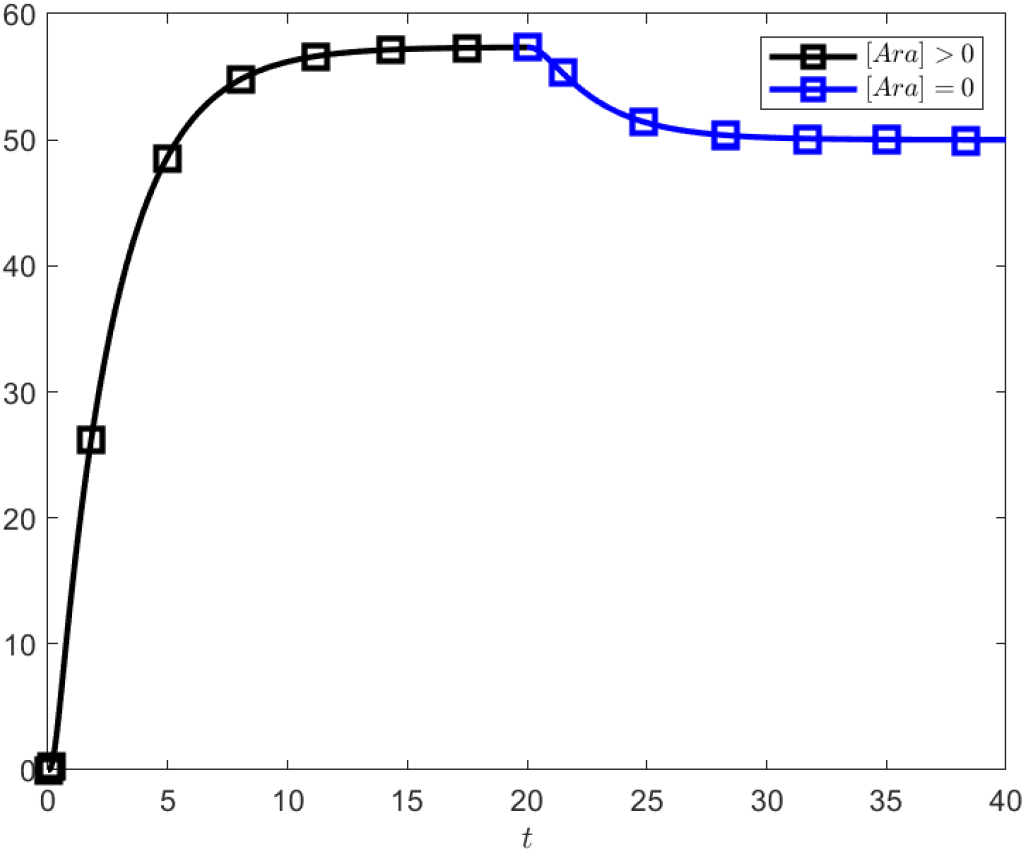
GFP protein concentration dependent of time varying arabinose concentration. This graph shows the circuits memory (hysteresis).

The parameter [*Ara*] *>* 0 is changed to [*Ara*] = 0 around the time unit ≈ 19, to mimic arabinose depletion. The system then seeks a new stable state where the GFP concentration diminishes but never becomes 0.

### F. Prediction of target protein concentration

It is necessary to be able to make predictions of the target protein (GFP) concentration for this circuit. Also knowing if this concentration is dependent with the initial concentration of inductor (arabinose). Since the equilibrium GFP concentration is the maximum we can achieve, the following case becomes of interest:

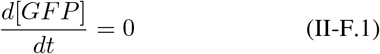

Substituting on II-B.6 and reorganizing

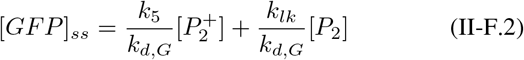

Where [*GFP*]_*ss*_is the equilibrium concentration of GFP. It is also known that 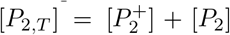, substituting and reorganizing.

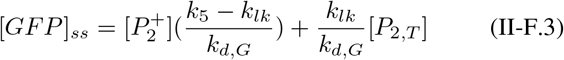

It becomes known here that the equilibrium concentration of GFP is a linear function of the activated promoter 2 concentration 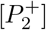. It is sufficient to establish which variables rule the equilibrium concentration. As studied in seccion II-C, this comes from:

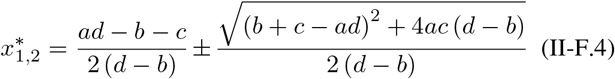

As studied, the 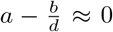 approximation will be used to prove the equilibriums stability. This approximation allows a simpler expression for *x*^*^.

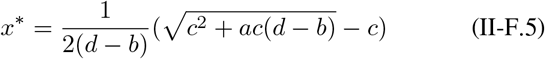

It is clear that *c* is an arabinose concentration related constant. Thus, both expressions can be graphed to get a notion of the relationship between *x*^*^ and *c* and hence, between [*GFP*]_*ss*_ and [*Ara*].(see fig. 9)

**Fig. 9.**
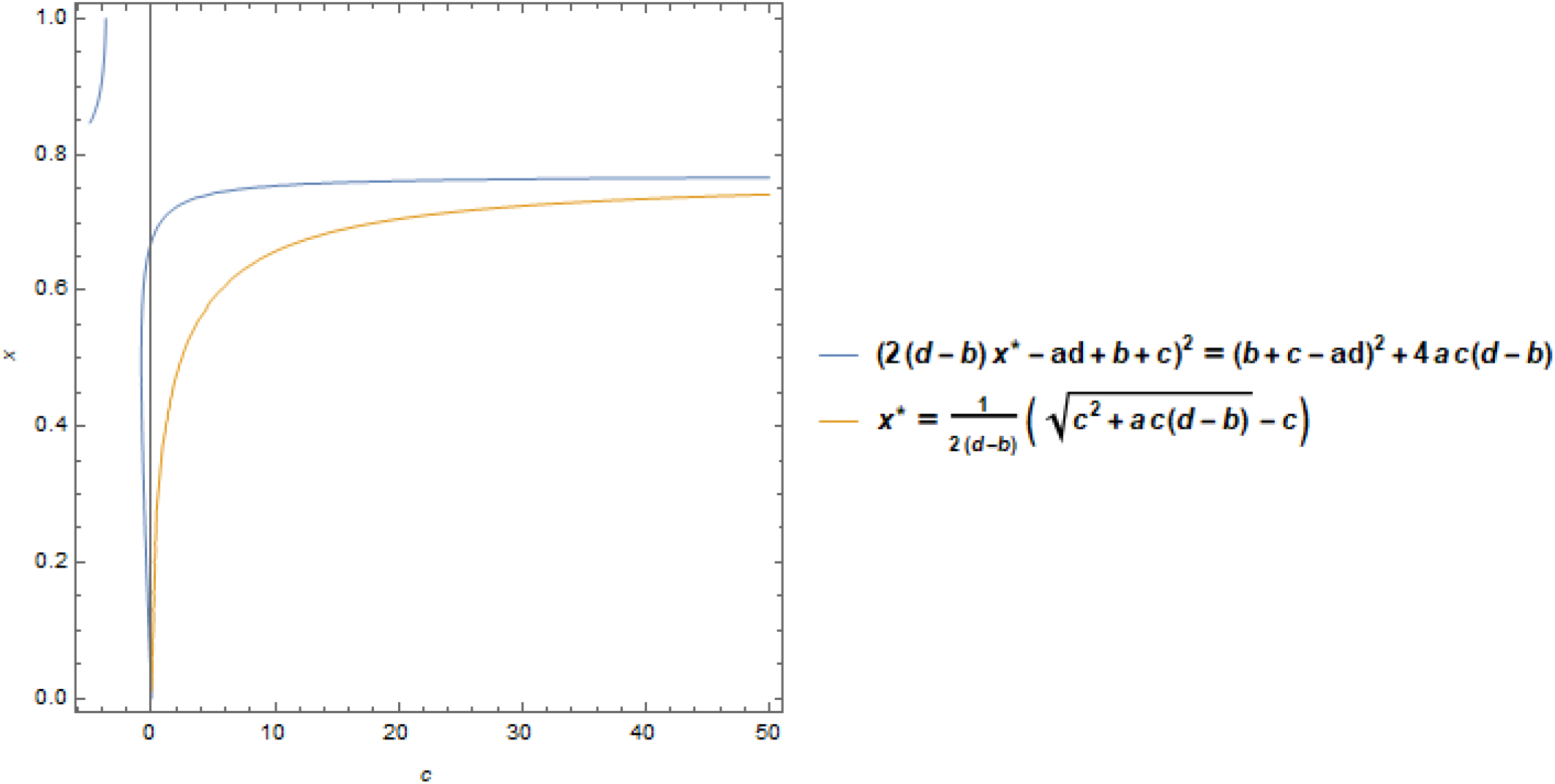
*x*^*^ dependency with *c*, reflecting [*GFP*]_*ss*_ dependency with [*Ara*].(*a* = 0.77, *b* = 1.1, *d* = 3.7)

Regardless of the chosen values, the behavior of *x*^*^ becomes rapidly independent of *c* while this value increases. This reinforces the hypothesis that the circuits behavior is nonlinear for arabinose, meaning, low arabinose levels can trigger high concentrations of interest protein. The circuit may behave in a “All or nothing” type of way regarding arabinose. Carrying out experiments becomes necessary to determine the minimum concentration of arabinose needed.

This allows the model to move away from the dependency that [*GFP*] and [*Ara*] may have, thus developing a more useful expression. Back to equation II-F.3,it is valid to assume that in equilibrium 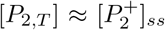, the equation is reduced to^6^

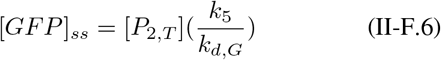

The plasmid concentration containing promoter 2 [*P*_2,*T*_] has been considered constant in this model. It can be assumed, in a different time scale, that it follows a Luedeking-Piret type kinetic.

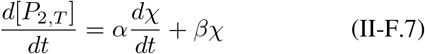

Where *χ* is cell concentration, *α* and *β* are constants of the model related with the synthesis of the metabolite. Plasmid production is solely associated with cellular growth, meaning *β* = 0, and consequently

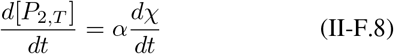

Integrating and substituting in II-F.6

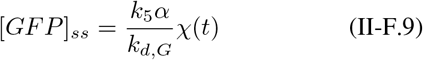

## III. Cellular automata model

### A. Model definition and simulation

The model to simulate consists of a genetic circuit inside a cell. GFP production (or any other targeted protein) with a low initiator intake is expected by this circuit using a positive feedback mechanism or self-initiator.

The system describes itself as a group of differential equations, but its answer can be hard to obtain. Advanced computation can only be approximate by a few decimals and show the system’s behavior. Thus, the making of a cellular automaton model.

Model’s characteristics are as described below:

Cellular space: Made by a square bi-dimensional matrix with repeating edge, meaning the final cells collide with initial cells of the new matrix, with variable size according to the experiment convenience, with each matrix side having 100 or 200 cells, each one having an element and number assigned. Cell status: Cells can be in either of the AC’s conditions described by Q=EV, PrAG, PrA, PAG, PA, PAC, Ar, AC, G, A1. Neighborhood: The neighborhood used in the model is the Moore neighborhood (see Fig.10), which involves orthogonal and diagonal neighbors. This type of neighborhood grants mobile individuals the ability to move in any direction, hence its use in this model.

**Fig. 10.**
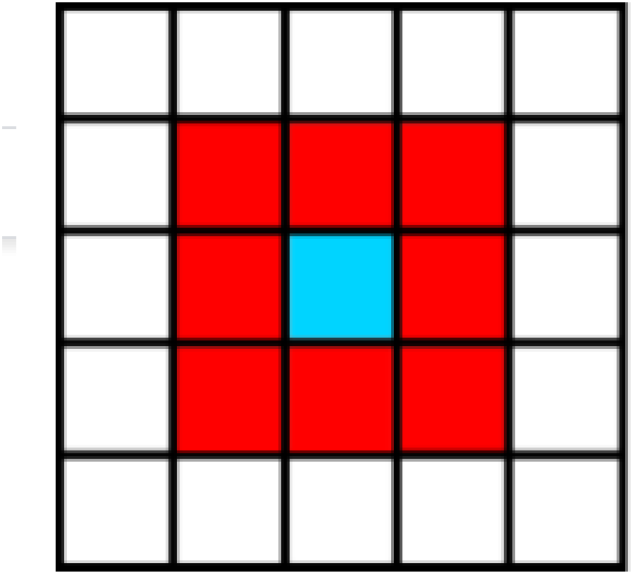
Neighborhood. Blue represents the active cell red depict their neighbors.

Figure 10. Initial arrangement: Before the simulation run, the location of promoters and producers inside the designated simulation area is fixed, which will not change over time. Run time: Whereas the experiment execution speed depends on the computer’s capability, the number of molecules produced or removed for every unit of time must concur with the rate in a real-time minute. Parameters: Selected parameters for the simulation run are: GFP protein production rate: 4 molecules per minute, AraC: 3 molecules per minute, Activator: 10 molecules per minute. Half-life: Half-life depends on the ongoing experiment.

Model evolution rules: Cells PrAG, PrA, PAG, PA, and PAC will not move through the simulation. EV can change locations with EV, Ar, AC, G, and A1. PrAG and PrA will constantly check for A1 and/or AC in its vicinities. PA cells will randomly turn EV cells into A1 cells at a rate of 10 cells per minute if PrA has either A1 or AC nearby. PAG will randomly turn EV cells into A1 and GFP at 10 and 4 cells per minute respectively if PrAG cell has an A1 cell next to it. PAC cell will randomly turn EV into AC at a rate of 3 cells per minute. Whilst Ar, Ac, G, and A1 will degrade according to their half-life. Collision control: As described in rule 2, only the EV cell can move through the matrix, allowing the space between cells to change, not vice versa, therefore preventing interaction between two molecules wanting to be in the same cell at the same time.

### B. Experimentation

For the testing phase, a code using Python programming language was written and executed in Google servers using the Google Colab platform, which does not have any graphic UI for any change made to the parameters will change the code itself. Apart from settling general system behavior, the model will additionally evaluate three different scenarios: Collection of activator protein: In the presence of glucose, arabinose molecules responsible for the activator production will not degrade. Therefore, activator protein will keep its output through arabinose and its self-induced mechanism, leading to hazardous arabinose pileup for the cell. It is worth mentioning that activator protein has not been widely analyzed yet, and its half-life is still uncertain. A half-life protein from bacteria will be used instead (roughly 20 hours). Self-activation faculty: The main feature of this circuit it’s the activator’s faculty of self-activation once arabinose depletion, hence the importance of checking the presence of this characteristic in the system. Broth composition could alter arabinose degradation. Still, usage of arabinose as the main carbohydrate for growth is not convenient due to high acquisition costs compared to other carbohydrates. For instance, Glucose in presence with arabinose will still be degraded, demonstrating the faculty of self-activation, nonetheless.

### C. Results and analysis

#### General system behavior

General system behavior shows the expected results, a regular non-accumulative AraC production, opposite to activator protein, for having two promoters’ induction has high concentration inside the cell.

#### Activator protein accumulation

The general system behavior (see Fig.11) shows high activator protein accumulation, although arabinose keeps degrading. A possible solution involves adding an amino acid sequence encouraging protein degradation, also known as degradation tale. Figure 12 shows the simulation with the degradation tale, which on average, demeanors arabinose half-life by half. Results lack approval since activator protein concentration is still very high and dangerous for the cell due to energy consumption or stress induced by such high concentrations. There are many other ways to reduce this concentration, such as exchanging the consensus sequence in promoters or RBS. These could potentially reduce the usage of resources given that reducing the half-life of the protein only increases degradation rates and keeps using the same amount of means for their synthesis. By changing the consensus sequences, the production rate of activator protein steadies, reducing its quantity and the resources used to produce them.

**Fig. 11.**
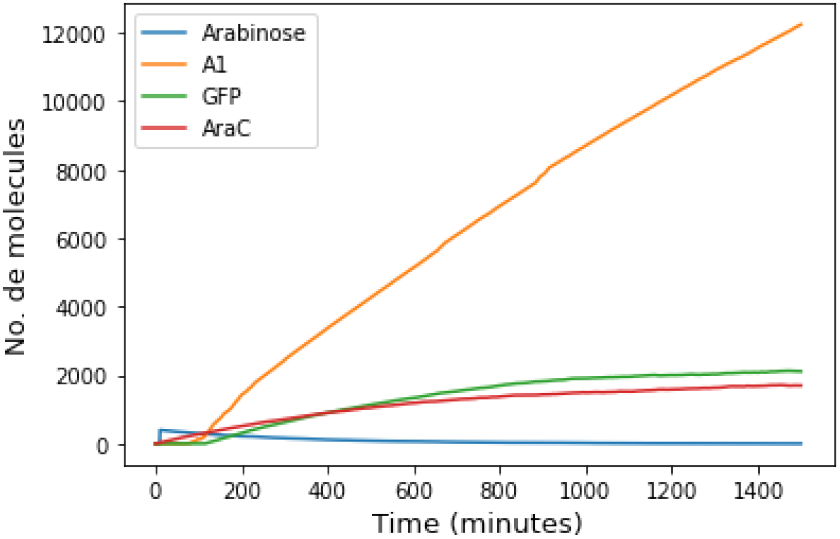
General behavior system simulation. Use of 200 cell side matrix, activator protein half-life of 1200 minutes, and arabinose half-life of 200 minutes.

**Fig. 12.**
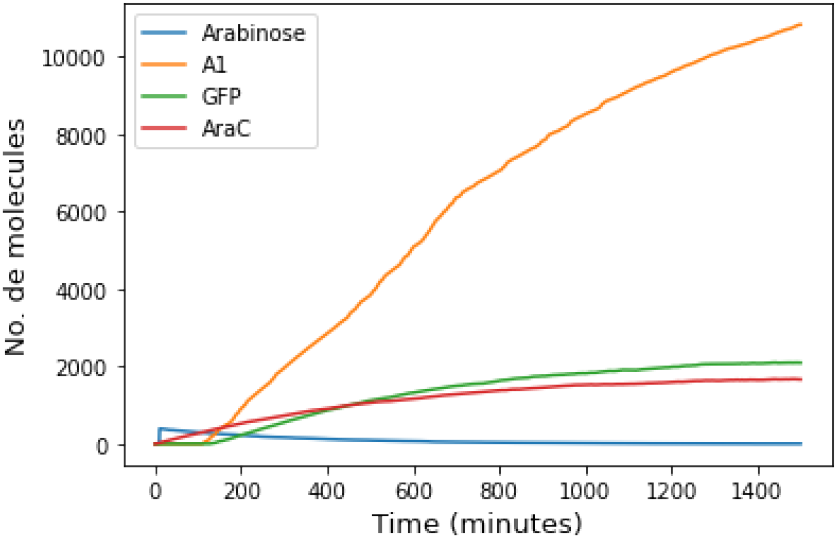
Half-life of activator protein has been cut in half (600 minutes) due to the addition of a degradation tale. However, it keeps a high concentration rate inside the cell.

Self-activation faculty To prove the self-activated mechanism’s effectiveness, even with a lower-than-expected half-life for the activator protein, a simulation with a low half-life for the activator protein and fast degradation rate of arabinose, as shown in figure 13, was run. At first, the simulation shows a rapid increase in activator protein due to arabinose induction and self-activation. As the arabinose decreases, the activator protein concentration also decreases until both reach a stable point. This value remains constant even after arabinose runs out.

**Fig. 13.**
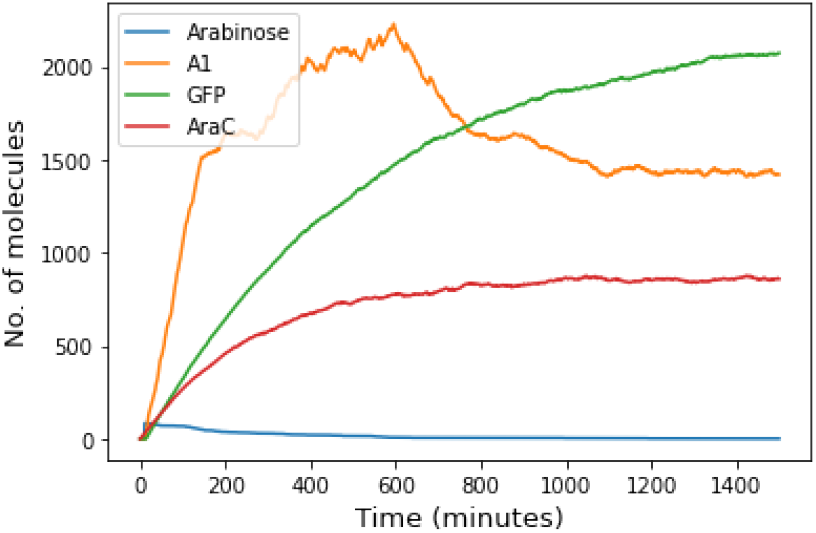
Decreasing the half-life of arabinose to 100 minutes and keeping activator protein half-life on 50 minutes shows the memory mechanism’s effectiveness.

## IV. Conclusions

### A. Mathematical model

Thanks to the proposed hypothesis (AraC and Arabinose in equilibrium), the differential equations model yields many interesting results. These will help predict the circuits real behavior. It is known now that the whole system dynamics can be understood by solely analyzing the dynamic between activator protein and activated promoter 2. This is due to the target protein (GFP) equilibrium concentration being a lineal function of the activated promoter 2 concentration 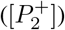. Therefore, the 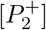 concentration is a reflex of the GFP concentration. Thanks to this a 2 equation non-dimensional system was obtained. Analyzing equilibrium states and via a justified approximation, it was proven that activator protein, 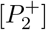 and GFP concentrations will arrive at a stable value in time while arabinose is present. This value will depend on system parameters (transcription factors bonding and dissociation force, protein degradation and production rate constants, plasmid concentration, arabinose concentration). At the same time this value will not be a function of its initial concentration presence of GFP, activator protein or 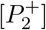 does not matter and the equilibrium concentration will always be reached. To exemplify this proven results, numeric solutions of activator protein, GFP and 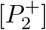 concentrations were run. It was observed that, initial conditions do not matter, and an equilibrium state will be reached. All the curves converge in a single value (see fig. 4, 5 y 6)

Equilibrium and system stability changes if there is no inductor presence and basal production is negligible. In this case, 2 equilibrium states were proven: First, when there is no GFP, activator and 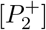 production. The system is in OFF state and the equilibrium is unstable. In the second equilibrium GFP, activator protein and 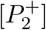 concentrations are positive and different from 0. This is the systems ON state, and it is stable. This can be summarized as:

- The system will be OFF exclusively if there is no arabinose, activator protein and 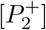 presence and basal production is negligible. Any fluctuation on the system can take it to its ON state.
- If the system is ON, it can not go back to an OFF state. Even after arabinose is completely exhausted, the system will maintain its ON state (hysteresis). This does not mean the ON states with and without arabinose have the same equilibrium values. The ON state without arabinose may produce less target protein, as shown in figure 8.

Finally, arguments were made against the search of a mathematical expression capable of predicting target protein (GFP) as an arabinose function. This is thanks to a graphic analysis of GFP versus arabinose concentration, indicating that dependence is quickly lost. Consequently, increasing arabinose concentration will rise that of GFP insignificantly. It seems that a minimum arabinose concentration yields a good GFP production. Nevertheless, it is possible that the time it takes for the system to reach equilibrium state its dependent con arabinose concentration. Experimentation for analyzing this time and the initial concentration of arabinose is needed to make any kind of correlation.

All this ends in a simple equation predicting target protein concentration in equilibrium in function of biomass concentration. Graphic analysis also identifies there is a strong correlation between target protein in equilibrium and activator protein kinetic stability. The more kinetically stable the activator protein is more target protein is produced. However, if the protein is too stable it will tend to accumulate inside the chassis causing stress. A search for the optimal stability that yields the maximum target protein concentration obtainable and the minimal stress possible is suggested.

### B. Cellular automata model

Despite the drawbacks produced by the uncertainty of the activator proteins half-life, GFP production is acceptable in every case. Is interesting to see that the production value is the same in all three situations, thus indicating self-activation is possible even at low activator protein concentrations. It becomes essential then, to know such proteins degradation rate via experimentation. This will allow to choose if a method to lower its concentration and the metabolic cellular stress is needed. Especially considering that the half life used in the model corresponds to that of the average protein and may no be a realistic value thanks to other factors not considered in the degradation rate simulation (e.g., protein size, sequences that may impact its degradation rate, etc.). Despite all this, simulations make it clear that self-activation with low arabinose quantities is possible, eliminating the need for other higher cost molecules like IPTG.

Visit https://github.com/OllinSynBioIPN/TETL-BOX to see all the code used in this work.

## V. Acknowledgments

We thank the Ollin SynBio IPN Student Research Group for all the support provided, also for wanting to make science accessible to everyone, and for the fantastic idea of producing reagents more economically. We also thank our advisor Claudia Hernandez for all the feedback provided, also for encouraging us to translate mathematical and computational results into biological results.

## Footnotes

1 In this work the following notation will be considered, let a transcription regulatory protein G, then G^+^ represents its conformation in which it can activate, G^−^ represents its conformation in which it can repress, now, let P be a regulatable promoter, then P^+^ represents the activated promoter, P^−^ represents the inhibited promoter and P represents the promoter in its basal state.

2 The constants used are just reaction rate constants obtained in the previous reactions.

3 The equation system non-dimensionalization is a long and technical process, and therefore has been omitted from this document.

4 Variable and non-dimensional constant’s meaning can be deducted from its definition: - *x* is the non-dimensional concentration of activated promoter 2. - *y* is the non-dimensional activator concentration. - *τ* is the non-dimensional time. - *a* is a measure of the activator protein kinetic stability. - *b* is a measure of the dissociation force between the activator protein and its promoter. - *c* is a measure of arabinose concentration and basal production rate. - *d* is a measure of the activated promoter 2 protein production.

5 In this work *x*^*^ and *y*^*^ indicate equilirium values, this means, 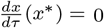 and 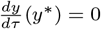 are true.

6 Assuming *k*_*lk*_ ≈ 0 leads to the same result.

